# Proteolytic Cleavage of the ENaC γ Subunit – Impact Upon Na^+^ and K^+^ Handling

**DOI:** 10.1101/2024.02.12.579964

**Authors:** Evan C. Ray, Andrew Nickerson, Shaohu Sheng, Rolando Carrisoza-Gaytan, Tracey Lam, Allison Marciszyn, Lei Zhang, Alexa Jordahl, Chunming Bi, Aaliyah Winfrey, Zhaohui Kou, Sebastien Gingras, Annet Kirabo, Lisa M. Satlin, Thomas R. Kleyman

## Abstract

The ENaC gamma subunit is essential for homeostasis of Na^+^, K^+^, and body fluid. Dual subunit cleavage before and after a short inhibitory tract allows dissociation of this tract, increasing channel open probability (P_O_), *in vitro*. Cleavage proximal to the tract occurs at a furin recognition sequence (^143^RKRR^146^ in mouse). Loss of furin-mediated cleavage prevents *in vitro* activation of the channel by proteolysis at distal sites. We hypothesized that ^143^RKRR^146^ mutation to ^143^QQQQ^146^ (^Q4^) in 129/Sv mice would reduce ENaC P_O_, impair flow-stimulated flux of Na^+^ (J_Na_) and K^+^ (J_K_) in perfused collecting ducts, reduce colonic amiloride-sensitive short circuit current (I_SC_), and impair Na^+^, K^+^, and body fluid homeostasis. Immunoblot of ^Q4/Q4^ mouse kidney lysates confirmed loss of a band consistent in size with the furin-cleaved proteolytic fragment. However, ^Q4/Q4^ male mice on a low Na^+^ diet did not exhibit altered ENaC P_O_ or flow-induced J_Na_, though flow-induced J_K_ modestly decreased. Colonic amiloride-sensitive I_SC_ in ^Q4/Q4^ mice was not altered. ^Q4/Q4^ males, but not females, exhibited mildly impaired fluid volume conservation when challenged with a low Na^+^ diet. Blood Na^+^ and K^+^ were unchanged on a regular, low Na^+^, or high K^+^ diet. These findings suggest that biochemical evidence of gamma subunit cleavage should not be used in isolation to evaluate ENaC activity. Further, factors independent of gamma subunit cleavage modulate channel P_O_ and the influence of ENaC on Na^+^, K^+^, and fluid volume homeostasis in 129/Sv mice, *in vivo*.

## INTRODUCTION

The epithelial Na^+^ channel (ENaC) mediates Na^+^ reabsorption in the aldosterone-sensitive distal nephron, respiratory tree, and distal colon (1-3). This channel is highly Na^+^-selective and is canonically comprised of three transmembrane subunits (α, β, and γ), each encoded by a gene that is critical for fluid and electrolyte handling (1, 2, 4). Post-translational proteolytic cleavage of ENaC modulates its activity. This was first suggested in studies demonstrating that short-circuit Na^+^ currents in toad bladders were inhibited by treatment with a serine protease inhibitor (5). Experimental application of the serine protease, trypsin, was subsequently shown to stimulate ENaC activity in *Xenopus* oocytes or in renal epithelial cells in culture (6). Evidence that extracellular proteases can enhance distal nephron Na^+^ reabsorption *in vivo* came from the observation that experimentally applied proteases stimulate ENaC activity in microdissected distal nephrons and in distal nephrons perfused with proteases via micropuncture (7, 8). In parallel, numerous studies show that ENaC subunits are processed by proteases *in vivo* and that this proteolytic processing is regulated by a variety of factors (7, 9, 10).

Both the α and γ subunits of ENaC are subject to proteolytic processing. Each of these two subunits contains a short inhibitory tract that suppresses channel open probability (P_O_) and can only be removed after two cleavage events occur: 1) at a site proximal (N-terminal) to the inhibitory tract, and 2) a site distal (C-terminal) to the tract (11). Removal of the α subunit’s inhibitory tract moderately stimulates the channel *in vitro*, compared to non-cleaved channels (12). Removal of the γ subunit’s inhibitory tract further and significantly increases channel activity, regardless of α subunit cleavage (13, 14). Based on these observations, we suggested that a loss of the γ subunit’s inhibitory tract has a dominant role in channel activation (14) Removal of the γ subunit inhibitory tract occurs following furin-mediated cleavage at a site proximal to its inhibitory tract (mouse γ^140^RKRR^143^) in combination with cleavage at any of a of sites just distal to the subunit’s inhibitory tract. Extracellular proteases capable of performing this second cleavage event include prostasin (also known as channel activating protease 1, or CAP1) (6, 15), TMPRSS2 (16), TMPRSS4 (CAP2) (17), neutrophil and pancreatic elastases (18), kallikrein (19), matriptase (CAP3) (20), cathepsins (21, 22), plasmin (23, 24) and urokinase-type plasminogen activator (25).

Numerous physiologic and pathophysiologic stimuli enhance proteolytic processing of the γ subunit. Body fluid volume depletion occurring in response to dietary Na^+^ deprivation enhances γ subunit cleavage in kidneys from rats and mice (9, 26-33). Increased dietary K^+^ increases the proteolytically processed fraction of the γ subunit in kidneys (34, 35). Aldosterone signaling in response to fluid depletion or hyperkalemia likely contributes to these findings, as aldosterone infusion in rats and mice stimulates γ subunit cleavage (9, 29, 36, 37), and primary aldosteronism in humans is associated with increased urinary protease expression that resolves following adrenalectomy (10). Proteolysis of the γ subunit is also increased in kidneys from animals with experimentally-induced proteinuria or humans with proteinuric kidney disease (23, 38-41). This phenomenon may occur as a consequence of leakage of plasma proteases into the tubule through damaged glomeruli. This suggestion is supported by the observation that proteases capable of activating ENaC are found in the urine of individuals with nephrotic syndrome (40), preeclampsia (42), and diabetic nephropathy (43, 44). These observations have led us and others to hypothesize that γ subunit cleavage contributes to Na^+^ retention in the setting of body fluid depletion (8), and proteinuric kidney disease (45-47).

Given the profound effects of ENaC γ subunit proteolysis noted with *in vitro* studies, we hypothesized that γ subunit proteolytic cleavage with the release of its inhibitory tract will impact extracellular volume and Na^+^ and K^+^ homeostasis. To test this hypothesis, we generated a mouse with genetic resistance to proteolytic removal of the γ subunit inhibitory tract. This was performed by replacing the γ subunit furin cleavage site (^140^RKRR^143^, or WT) with residues conferring resistance to furin cleavage (^140^QQQQ^143^, or Q4). This mutation successfully prevented activation of mouse ENaC by prostasin in *Xenopus* oocytes. We confirmed the viability of homozygous Q4 (γ^Q4/Q4^) mice and determined biochemically that this mutation prevented proteolytic processing at the furin cleavage site of the γ subunit. We examined the P_O_ of ENaC using on-cell patch clamp of principal cells in isolated connecting tubule/collecting ducts (CNT/CCDs) and measured transepithelial Na^+^ and K^+^ fluxes in microperfused tubules. We measured whole blood electrolytes including K^+^ and tCO_2_, and plasma aldosterone in mice on low Na^+^ and high K^+^ diets. We examined amiloride-sensitive currents in the distal colon. We assessed whole body fluid volume in response to low Na^+^ diet using live animal quantitative magnetic resonance. Surprisingly, the phenotypic differences between the WT and Q4 mice were modest, suggesting that other factors have important roles in regulating ENaC activity.

## MATERIALS AND METHODS

### Two-electrode voltage clamp electrophysiology

cDNA encoding the mouse γ subunit was mutated to replace ^140^RKRR^143^ with ^140^QQQQ^143^ using the QuckChange II XL mutagenesis kit (Agilent Technologies). cDNAs encoding ENaC’s mouse α, β, and γ subunits and mouse prostasin were transcribed using the mMessage mMachine transcription kit (Life Technologies). Resulting cRNA encoding each of the subunits was injected into stage V to VI *Xenopus* oocytes. Oocytes were incubated in Barth’s saline for 24 to 48 hours to allow expression of the channel. Whole cell currents were recorded using standard two-electrode voltage-clamp techniques as previously described (48, 49).

### Generation of mouse models

All animal experiments were approved by the Institutional Animal Care and Use Committee at the University of Pittsburgh or at the Icahn School of Medicine at Mt. Sinai, where appropriate. Animals were housed in AAALAC-accredited facilities at respective institutions. Mice in the 129/Sv background (129S2/SvPasCrl, Charles River Laboratories) were genetically modified using CRISPR-Cas9 technology, as previously reported (50). Exon 3 of the *Scnn1g* locus on mouse chromosome 7 was modified using a Cas9 sgRNA targeting the sequence CCTCGGAAACGCCGGGAAGCAGG, changing amino acids ^140^RKRR^143^ in the ENaC g subunit to ^140^QQQQ^143^ and to add a BseY1 restriction endonuclease target sequence (CCCAGC) to facilitate genotyping of genetically modified animals. Genotyping was performed by polymerase chain reaction-based amplification of the surrounding region of *Scnn1g* with primers ER61 (GACTGTGGGACTACCAGCTC) and ER67 (AAGGGACTGGTCAGGAGACA), resulting in a 498 nucleotide product. Restriction endonuclease digestion with BseY1 resulted in 213 and 285 nucleotide DNA bands in ^140^QQQQ^143^, but not littermate control, mice. Mice were housed in facilities with a 12 hr/12 hr light/dark cycle. Standard mouse chow for breeding and growth was Prolab^®^ Isopro^®^ RMH 3000 (0.23% Na^+^, 0.94% K; LabDiet). Low Na^+^ diet was Teklad TD.90228 (0.01-0.02% Na^+^, 0.8% K^+^; Envigo). High K^+^ diet was Teklad TD.09075 (0.3% Na^+^, 5.2% K^+^ as KCl). All mice were provided with reverse osmosis-purified drinking water, *ad libitum*. Mice were sacrificed under general anesthesia provided by inhaled isoflurane.

### Mouse body composition measurements

Body composition of mice was measured in live, unanesthetized 12 week-old mice by quantitative magnetic resonance using a 100H Body Composition Analyzer (EchoMRI) as previously described (51-53). Baseline body weight and body composition were measured daily for three days. On the third day, mice were transitioned to a low Na^+^ diet. Body weight and body composition was measured for 10 days on the low Na^+^ diet. All data shown are normalized to each animal’s average baseline body weight or composition.

### Mouse blood metabolite measurement

Blood was collected by intracardiac aspiration at the time of sacrifice from mice 10 to 16 weeks of age. Electrolytes were measured immediately on whole blood using an iSTAT analyzer (Abbott). Measurement of plasma aldosterone was accomplished by separating plasma from red blood cells in heparinized plasma-separator tubes, freezing in liquid nitrogen, and storing at -80 °C until analysis. Plasma was then thawed on ice, and aldosterone levels were measured using and aldosterone ELISA kit (Enzo).

### ENaC γ subunit immunoblots

Immunoblots were prepared from kidneys collected at the time of sacrifice from 12 week-old animals provided with a low Na^+^ diet for 10 days. Kidneys were snap frozen in liquid nitrogen and stored at -80°C until use. Once thawed, whole kidneys were homogenized in CelLytic MT lysis buffer (Sigma #C3228) at 1:20 weight/volume using a glass homogenizer. Lysates were then centrifuged at 12,000 x g for 20 minutes to remove insoluble material. After determining sample protein concentration by BCA assay, 20 mg of protein from each sample was subjected to PNGase F treatment (NEB #P0704) to remove N-linked oligosaccharides prior to SDS-PAGE. This technique allows for greater separation of distinct γ-ENaC cleavage products detectable by immunoblot (54). Proteins were resolved via SDS-PAGE using 4-15% Tris-glycine gels (Biorad) and subsequently transferred to PVDF membranes. After transferring, membranes were blocked in 5% milk dissolved in Tris-buffered saline with 0.1% Tween-20 (TBST) for 2 hours at room temperature, followed by overnight incubation in blocking buffer containing a primary antibody to the C-terminal region of γ-ENaC (StressMarq #SPC-405; 1:1,000 dilution). On the following day, membranes were washed 5 times in TBST before incubating in blocking buffer containing HRP-conjugated secondary antibody for 1 hour at room temperature. After washing again 5 times with TBST, chemiluminescent substrate (Biorad Clarity; #1705060) was applied and membranes were imaged with a Chemidoc multipurpose imager. Densitometric quantitation of γ-ENaC cleavage product abundance relative to the full-length protein was determined using ImageJ software.

### Connecting tubule/collecting duct single channel recording

Patch clamp studies were performed on manually dissected tubules from kidneys harvested from 14 to 17 week-old mice provided with a low Na^+^ diet for six to seven days. The mid-section of the mouse kidney was sliced into 1 mm sections with a sharp razor, from which the renal cortex was removed and cut into small blocks. The blocks were then treated with 1 mg/ml collagenase type II in Leibovitz’s L-15 Medium (Thermo Fisher Scientific Inc.) at 37°C for 30-45 minutes. The cortex blocks were washed with cold L-15 media and dissected immediately or stored temporarily on ice. Connecting tubules/cortical collecting ducts (CNTs/CCDs) were isolated manually with dissecting needle and tweezers and transferred to a cover glass coated with poly-lysine (Sigma-Aldrich, Inc., St. Louis, MO) for patch clamp studies, as previously described (55). CNTs/CCDs were opened with sharp pipettes under an inverted microscope, and principal cells were identified by morphology. Heat-polished patch pipettes were filled with a pipette solution containing 140 mM LiCl, 2 mM MgCl_2_, and 10 mM HEPES at pH 7.40 with a tip resistance of 4-7 MΩ. Cell-attached patch clamp experiments were conducted at room temperature using a PC-ONE Patch Clamp Amplifier (Dagan Corporation, Minneapolis, MN), DigiData 1440A, and Clampex 10.4 software (Molecular Devices, San Jose, CA). Patches were clamped at multiple voltages to obtain current-voltage (IV) relationships, and long recordings (>5 min) were performed at -60 mV to determine open probability. Data were sampled at 10 kHz and filtered at 1 kHz. Single-channel recording data were analyzed using Clampfit 10.4. Single-channel activity (NP_O_) was obtained using the single-channel searching function of Clampfit 10.4 (Molecular Devices), and channel number was determined by visual inspection of the whole recordings. Unitary currents were estimated by cursor measurements and represented by the average of three measurements at a clamping voltage. The slope conductance was estimated from the linear fit of unitary currents and clamping voltages in the range of -20 to -100 mV.

### Colonic epithelium electrophysiology

Mice were placed on a low Na^+^ diet for 14 days and sacrificed at an age of 10 to 16 weeks for harvesting of colons. Mucosal/submucosal colon preparations were prepared by opening the colon longitudinally along the mesenteric line, pinning the tissue mucosal side down in a dissecting dish, and peeling away the muscularis layers using fine forceps. Tissues were then mounted on 0.3 cm^2^ sliders for use in an Ussing-style recording chamber (Physiologic Instruments) and bathed in Ringer’s solution containing, in mM: 140 Na^+^, 119.8 Cl^-^, 25 HCO ^-^, 5.2 K^+^, 1.2 Ca^2+^, 1.2 Mg^2+^, 2.4 HPO_4_ ^2-^, 0.4 H PO ^-^, and 10 glucose. The solution was warmed to 37°C and maintained at pH 7.41 by gassing with 5% CO_2_ balanced with oxygen. Tetrodotoxin (0.5 μM) was added to the serosal bath to inhibit neurogenic secretion. After allowing approximately 20 minutes for equilibration, short-circuit current (I_SC_) was measured using a multichannel voltage clamp/amplifier (VCC MC6; Physiologic Instruments) controlled by computer-operated software (pClamp 10, Molecular Devices). After currents stabilized, ENaC-dependent Na^+^ absorption was measured as the change in I_SC_ after addition of 100 μM amiloride to the apical bath compartment.

### Isolation and in vitro microperfusion of CCDs for measurement of net transepithelial Na^+^ and K^+^ transport

Tubular Na^+^ and K^+^ fluxes (J_Na_ and J_K_, respectively) were measured in isolated, perfused collecting ducts, as previously described (56-58). Eight- to 10-week-old mice were fed a low Na^+^ diet for six to seven days before sacrifice, at which time kidneys were harvested, sectioned coronally, and placed in chilled Ringer’s solution (containing, in mM: 145 NaCl, 2.5 K_2_HPO_4_, 2.0 CaCl_2_, 1.2 MgSO_4_, 4.0 Na^+^ lactate, 1.0 Na^+^ citrate, 6.0 L-alanine, and 5.5 glucose. pH 7.4). Single collecting ducts(0.3 to 0.4 mm in length) were manually microdissected and transferred to a temperature and O_2_/CO_2_ controlled specimen chamber, mounted on concentric glass pipettes, and perfused and bathed at 37°C with Burg’s solution containing, in mM: 120 NaCl, 25 NaHCO_3_, 2.5 K_2_HPO_4_, 2.0 CaCl_2_, 1.2 MgSO_4_, 4.0 Na^+^ lactate, 1.0 Na_3_ citrate, 6.0 L-alanine, and 5.5 D-glucose; pH 7.4. During 45 min of equilibrium and thereafter, the bath solution was suffused with 95% O_2_ and 5% CO_2_ and continuously replaced at a rate of 10 mL/hour using a syringe pump (Razel Scientific). After equilibration, three to four samples of tubular fluid per experimental condition were collected under water saturated light mineral oil by timed filling of a calibrated ∼7 nL volumetric constriction pipette. Each tubule was perfused at low (∼1.0 ± 0.1 nL•min^-1^•mm^-1^) and high (∼5.0 ± 0.2 nL•min^-1^•mm^-1^) flow rates. The flow rate was varied by adjusting the height of the perfusate reservoir. At the end of each experiment, ouabain (200 mM) was added to the bath to inhibit all active transport, and three additional samples of tubular fluid were obtained for analysis to determine the composition of the solution actually delivered to the lumen of the tubule. Na^+^ and K^+^ concentrations of perfusate and collected tubular fluid were determined by helium glow photometry. The rates of net ion transport (J_X_, in pmol•min^-1^•mm^-1^), were calculated using previously described standard flux equations (59). As transport measurements were performed in absence of transepithelial osmotic gradients, water transport was assumed to be zero. Calculated ion fluxes were averaged to obtain a mean rate of ion transport for the CCD at each flow rate.

### Statistics

All statistics were performed using GraphPad Prism v. 9.5.0. Outlier testing was performed using the ROUT method (Q = 1%). Pair-wise comparisons were performed using a two-sided Student’s *t*-test with α < 0.05 considered significant. For time series analysis (body composition as a function of time and genotype after transition to a low Na^+^ diet), significance was determined by mixed-effects analysis.

## RESULTS

To confirm that disruption of the furin cleavage site (^140^RKRR^143^, WT) in the mouse ENaC γ subunit impairs proteolytic activation of the channel, we mutated the furin cleavage site (to ^140^QQQQ^143^, Q4) in the cDNA encoding the γ subunit and examined the influence of the extracellular protease, prostasin, on ENaC currents in *Xenopus* oocytes. RNA made from the cDNAs of mouse α, β, and γ subunits was injected into oocytes with, or without RNA encoding mouse prostasin (**Figure 1**). Measured currents were confirmed to be ENaC currents by perfusion of the oocyte with the ENaC blocker, amiloride at a concentration of 10 µM. Amiloride-sensitive current magnitudes were significantly greater in oocytes co-expressing prostasin with WT ENaC than in oocytes expressing ENaC alone. In contrast, currents in oocytes with Q4 ENaC and co-expressing prostasin were no different than in oocytes expressing the Q4 subunit with no prostasin. These data confirmed that substitution of ^140^RKRR^143^ with ^140^QQQQ^143^ rendered the mouse ENaC channel insensitive to activation by an extracellular serine protease.

**Figure 1.**
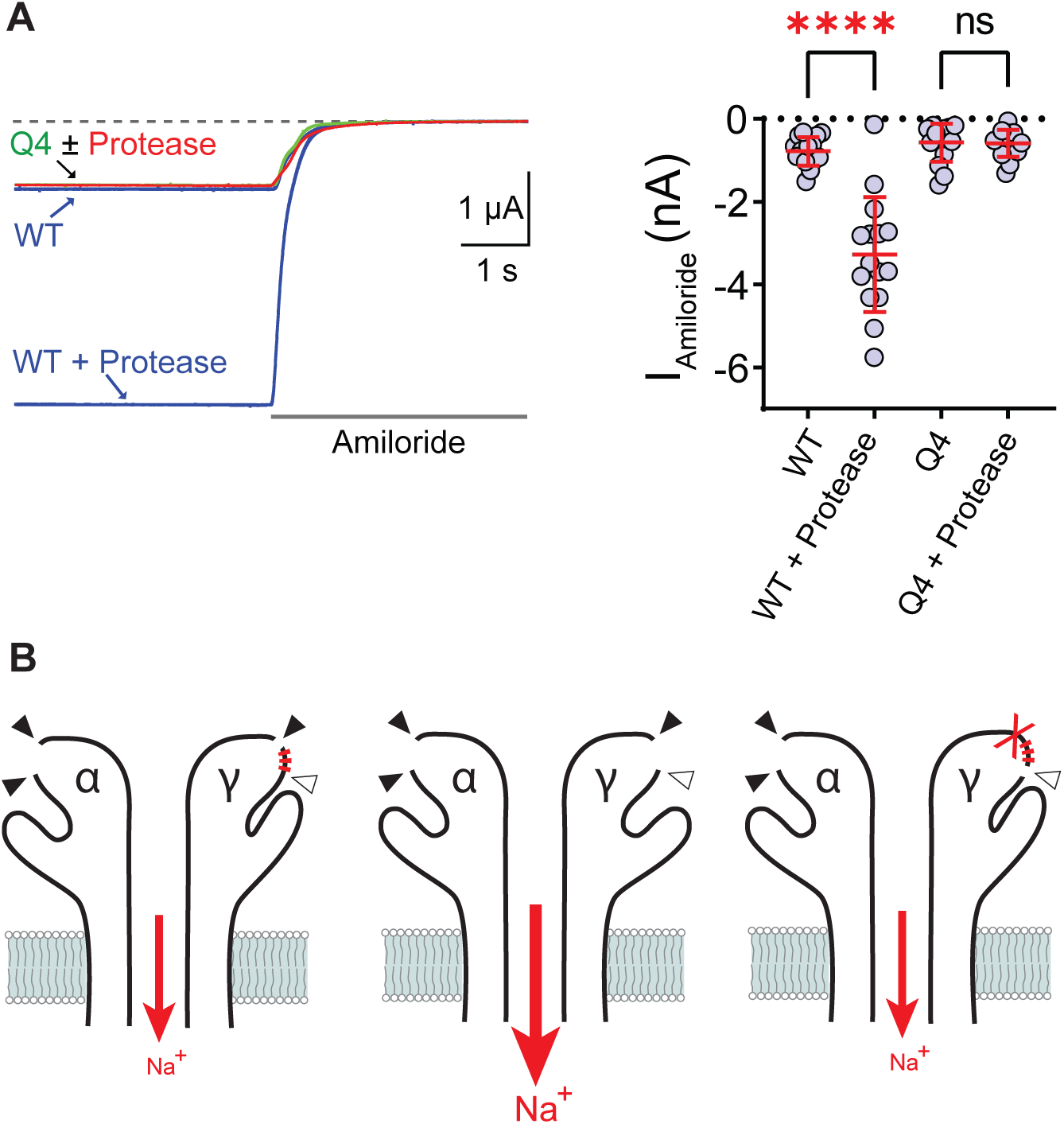
Disruption of the mouse ENaC γ subunit furin cleavage site, ^140^RKRR^143^ (WT), by replacement with ^140^QQQQ^143^ (Q4) impairs proteolytic activation of the channel. A: Representative two-electrode voltage clamp current recordings are shown (left) from *Xenopus* oocytes expressing mouse ENaC α and β, subunits and a γ subunit that is either WT or Q4. ENaC was co-expressed with or without the serine protease prostasin. Current amplitudes are shown (right). Na^+^ currents were similar in WT and Q4 ENaC in the absence of prostasin. Co-expression of WT, but not Q4, ENaC dramatically increases current amplitude (N = 15 oocytes per group, **** p < 0.001, ANOVA). Error bars represent standard deviation of the mean. B: Schematic showing proteolytic activation of ENaC. For clarity, only α and γ subunits are depicted. Cleavage of the α subunit at furin recognition sequences (solid arrowheads) removes an inhibitory tract (not shown), resulting in a channel with intermediate open probability (P_O_) (left). Removal of the γ subunit’s inhibitory tract (red hatch marks) requires cleavage at both a furin recognition sequence proximal to the tract *and* at a site distal to the tract (open arrowheads). Dual cleavage of the γ subunit results in loss of the γ subunit inhibitory tract, producing a highly active channel with a high P_O_ (middle). Disruption of the γ subunit’s furin recognition sequence (red X, indicating the Q4 substitution) abrogates activation of the channel by proteases that cleave distal to the γ subunit inhibitory tract (right).

Next, we examined how the loss of proteolytic activation of ENaC influences *in vivo* electrolyte handling. Using a CRISPR-Cas9 strategy in mice, we modified *Scnn1g*, encoding ENaC’s γ subunit, to reproduce the γ subunit ^140^QQQQ^143^ mutation that abrogated proteolytic activation of ENaC by prostasin in *Xenopus* oocytes (**Figure 2**.) Mice were bred into the 129/Sv background. Mice lacking expression of the ENaC γ subunit exhibit perinatal mortality (1). However, there was no evidence of perinatal mortality in γ^Q4/Q4^ mutant mice, as crosses of heterozygous parent mice resulted in litters with γ^WT/WT^, γ^WT/Q4^, and γ^Q4/Q4^ pups in ratios consistent with Mendelian inheritance (**Table 1**). Health and behavior of γ^Q4/Q4^ mice were subjectively normal. Weights of adult γ^Q4/Q4^ mutant mice resembled γ^WT/WT^ littermate controls (**Table 2**). Homozygous male γ^Q4/Q4^ mice were fertile.

**Figure 2.**
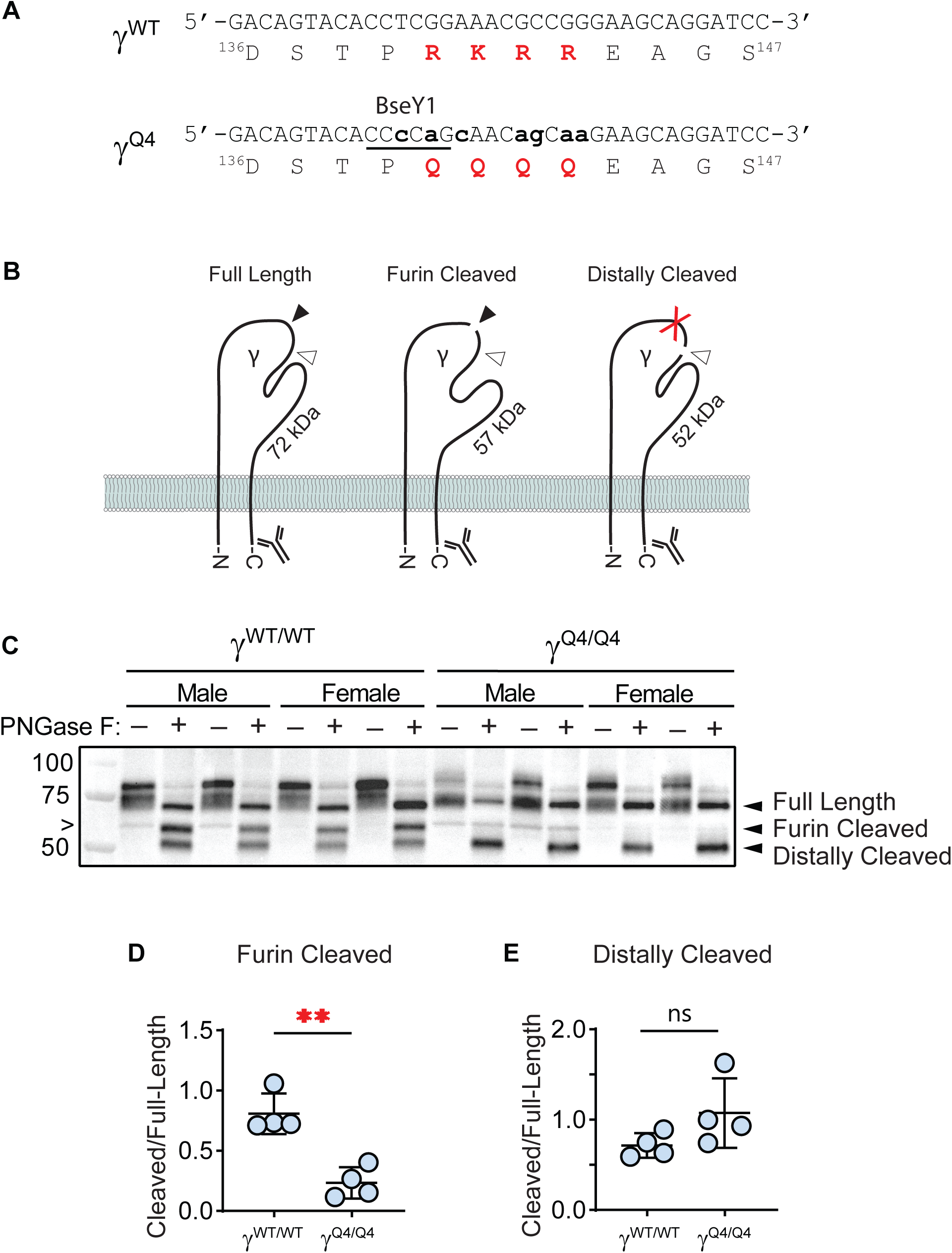
γ^Q4/Q4^ mice demonstrate impaired γ subunit proteolytic processing. A: DNA and amino acid sequences within the mouse *Scnn1g* locus and the ENaC γ subunit. Emboldened, lower case nucleosides represent those modified by CRISPR/Cas9. Underlined nucleosides indicate a silent BseY1 restriction endonuclease sequence introduced to identify genetically modified mice. Red amino acids represent the wild-type or disrupted γ subunit furin recognition sequence. B: Schematic of the anticipated effects of the Q4 substitution on the γ subunit cleavage fragments as detected by immunoblot using a C-terminal anti-γ subunit antibody. Full-length γ subunit following PNGase F treatment migrates at 72 kDa. Cleavage of the γ subunit at the furin recognition sequence (filled arrowhead) results in a 57 kDa C-terminal fragment. Cleavage distal to the inhibitory tract (open arrowhead) results in a 52 kDa cleavage fragment. Mutation of the furin recognition sequence with Q4 (red X) would be predicted to eliminate the 57 kDa cleavage fragment. C: Immunoblot of kidney lysates from γ^Q4/Q4^ and γ^WT/WT^ control mice. Immunoblots of lysates run before (-) or after (+) treatment with PGNase F, used to remove N-linked oligosaccharides from the subunit, improving resolution of the proteolytic cleavage fragments. “>” notes a non-specific band, predominantly in kidney from male mice, of unclear significance. D: Combined quantification of 57 kDa band intensity from males and females, indicative of the furin-cleaved γ subunit, normalized to full-length, 72 kDa band intensity, reveals significant reduction of the 57 kDa band (N=4 γ^WT/WT^ and 4 γ^Q4/Q4^ mice including 2 females and 2 males in per group. **: p < 0.01 as determined by unpaired *t*-test). Our quantification included the non-specific band the migrated close to the 57 kDa C-terminal furin-cleavage fragment. E: Quantification of 52 kDa band intensity, normalized to full-length, 72 kDa band intensity, reveals no significant difference in γ^Q4/Q4^ as compared with γ^WT/WT^ mice (p = NS, unpaired t-test). Error bars represent standard deviation of the mean.

**Table 1.** Genotypes of pups from crosses of heterozygous (γ^WT/Q4^) mice.

| | $\gamma^{WT/WT}$ | $\gamma^{WT/Q4}$ | $\gamma^{Q4/Q4}$ |
| --- | --- | --- | --- |
| **Number of pups** | 87 | 183 | 89 |
| **% Pups per Litter** | $24 \pm 15 \%$ | $53 \pm 22 \%$ | $24 \pm 16 \%$ |

N = 59 litters. Errors represent standard deviation of the mean.

**Table 2.** Adult mouse weights.

| | $\gamma^{WT/WT}$ | $\gamma^{Q4/Q4}$ |
| --- | --- | --- |
| **Males** | $25.6 \pm 2.7 \text{ g (11)}$ | $25.9 \pm 2.4 \text{ g (11)}$ |
| **Females** | $23.0 \pm 2.4 \text{ g (20)}$ | $21.7 \pm 2.3 \text{ g (14)}$ |

Weights of mice ages 10 to 16 weeks. P = NS for within-sex comparisons by Student's *t*-test. Errors represent standard deviation of the mean.

To confirm that the γ subunit ^140^RKRR^143^ to ^140^QQQQ^143^ mutation reduced production of the γ subunit furin cleavage product, we placed mice on a low (< 0.02%) Na^+^ diet for 10 days and harvested kidneys. Dietary Na^+^ depletion enhances proteolytic processing of the γ subunit in an aldosterone-dependent fashion (9, 60). As seen in **Figure 2**, immunoblots of kidney lysates from γ^WT/WT^ mice probed with a C-terminal γ subunit antibody exhibited three predominant bands following PNGase F treatment to separate the γ-ENaC cleavage products. The full-length γ subunit migrated with an apparent molecular weight of 72 kDa. A 52 kDa band is consistent with cleavage distal to the channel’s inhibitory tract. A 57 kDa band is consistent with the C-terminal γ subunit furin cleavage product. In lysates from γ^WT/WT^ kidneys, the furin cleavage product / full length subunit abundance was high (0.80 ± 0.17, N = 4). However, in lysates from γ^Q4/Q4^ kidneys, the furin cleavage product / full length subunit abundance was significantly lower (0.24 ± 0.13, N = 4; *p* = 0.002), suggesting reduced proteolytic processing by furin. The extent of furin cleavage of γ^Q4/Q4^ is likely much lower, as we observed a non-specific band (noted with “>”) in males migrating at a slightly higher molecular weight than the γ furin cleavage product following PNGase F treatment. This band was present in the absence and presence of PNGase F treatment in the male γ^Q4/Q4^ lanes. In contrast, the distal cleavage product / full length subunit abundance did not differ significantly between γ^WT/WT^ and γ^Q4/Q4^ kidneys (0.72 ± 0.14, N = 4 for γ^WT/WT^, 1.1 ± 0.39, N = 4 for γ^Q4/Q4^; *p* = NS), indicating that processing by other proteases was not altered.

We examined blood electrolytes in γ^WT/WT^ and γ^Q4/Q4^ mice. In mice mice on a standard (0.23% Na^+^, 0.94% K^+^) diet, we observed no differences in blood Na^+^, K^+^, Cl^-^, total carbon dioxide (tCO_2_), blood urea nitrogen (BUN), hemoglobin (Hb), or ionized calcium (iCa^2+^) (**Table 3**). As mice with large reductions in expression of specific ENaC subunits exhibit increased sensitivity to dietary Na^+^-restriction (61), we examined plasma electrolytes and aldosterone levels in response to a low Na^+^ (0.01-0.02% Na^+^, 0.8% K^+^) diet. No differences in plasma electrolytes or aldosterone levels were observed. ENaC activity is largely required for connecting tubule and collecting duct K^+^ secretion, although a component of ENaC independent K^+^ secretion has been observed in rodents on a high K^+^ diet (62). We examined plasma electrolytes in mice on a high K^+^ diet (0.3% Na^+^, 5.2% K^+^ as KCl). With the exception of a higher plasma Cl^-^ concentration in male γ^Q4/Q4^ mice, no differences in plasma electrolytes were observed on this diet.

**Table 3.** Mouse Blood Electrolytes.

|  | Male | Male | Female | Female |
| --- | --- | --- | --- | --- |
| | $\gamma^{WT/WT}$ | $\gamma^{Q4/Q4}$ | $\gamma^{WT/WT}$ | $\gamma^{Q4/Q4}$ |
| **Standard Diet** |  |  |  |  |
| **Mice (N)** | 9 | 7 | 12 | 7 |
| **Na<sup>+</sup>** | 146 ± 1 | 146 ± 2 | 145 ± 2 | 145 ± 1 |
| **K<sup>+</sup>** | 4.3 ± 0.2 | 4.4 ± 0.6 | 4.1 ± 0.6 | 4.3 ± 0.4 |
| **Cl<sup>-</sup>** | 113 ± 3 | 113 ± 2 | 113 ± 2 | 113 ± 3 |
| **tCO<sub>2</sub>** | 23 ± 2 | 23 ± 2 | 22 ± 3 | 23 ± 2 |
| **BUN** | 29 ± 3 | 31 ± 4 | 30 ± 5 | 28 ± 7 |
| **Hb** | 13.9 ± 0.6 | 13.8 ± 0.9 | 13.3 ± 1.0 | 13.6 ± 1.1 |
| **iCa<sup>2+</sup>** | 1.27 ± 0.08 | 1.24 ± 0.09 | 1.30 ± 0.07 | 1.34 ± 0.06 |
| **Low Na<sup>+</sup> Diet** |  |  |  |  |
| **Mice (N)** | 7 | 7 | 8 | 7 |
| **Na<sup>+</sup>** | 146 ± 1 | 147 ± 2 | 147 ± 2 | 148 ± 2 |
| **K<sup>+</sup>** | 4.7 ± 0.4 | 4.2 ± 0.6 | 4.2 ± 0.2 | 4.7 ± 0.3 |
| **Cl<sup>-</sup>** | 117 ± 2 | 116 ± 3 | 118 ± 2 | 118 ± 1 |
| **tCO<sub>2</sub>** | 21 ± 2 | 21 ± 1 | 21 ± 2 | 21 ± 2 |
| **BUN** | 28 ± 5 | 26 ± 2 | 25 ± 5 | 26 ± 4 |
| **Hb** | 14.1 ± 0.3 | 14.3 ± 0.4 | 14.1 ± 0.6 | 14.6 ± 0.9 |
| **iCa<sup>2+</sup>** | 1.34 ± 0.04 | 1.31 ± 0.05 | 1.32 ± 0.05 | 1.33 ± 0.07 |
| **Aldosterone<sup>a</sup>** | 787 ± 189 | 664 ± 151 | 789 ± 134 | 745 ± 256 |
| **High K<sup>+</sup> Diet** |  |  |  |  |
| **Mice (N)** | 10 | 7 | 13 | 14 |
| **Na<sup>+</sup>** | 147 ± 2 | 148 ± 1 | 147 ± 3 | 146 ± 2 |
| **K<sup>+</sup>** | 5.5 ± 1.0 | 6.1 ± 0.9 | 6.3 ± 0.7 | 6.6 ± 0.7 |
| **Cl<sup>-</sup>** | 120 ± 5 | 124 ± 1<sup>b</sup> | 121 ± 4 | 121 ± 4 |
| **tCO<sub>2</sub>** | 21 ± 2.2 | 19 ± 1.6 | 19 ± 2.2 | 20 ± 0.9 |
| **BUN** | 25 ± 3 | 25 ± 3 | 25 ± 5 | 27 ± 6 |
| **Hb** | 13 ± 1 | 13 ± 1 | 13 ± 1 | 13 ± 1 |
| **iCa<sup>2+</sup>** | 1.32 ± 0.03 | 1.32 ± 0.08 | 1.31 ± 0.06 | 1.34 ± 0.08 |

<sup>a</sup>N for aldosterone measurements = 5 per group. <sup>b</sup>P < 0.05 for comparison with same-sex $\gamma^{WT/WT}$ littermate controls, by Student's *t*-test. Errors represent standard deviation of the mean.

Mice with reduced or absent expression of ENaC subunits exhibit impaired body fluid retention and increased sensitivity to a low Na^+^ diet (1, 61, 63). We therefore asked whether γ^Q4/Q4^ mice exhibit increased body fluid loss on a low Na^+^ diet. Male or female mice underwent measurement of body weight and total body water via quantitative magnetic resonance. Baseline measurements were performed for three consecutive days on a standard Na^+^ diet before mice were transitioned to the low Na^+^ diet. Measurements were then performed daily for a total of 10 measurements after transition to the low Na^+^ diet. Values were normalized to standard diet measurements and plotted in **Figure 3**. Mouse weights declined only slightly in response to the low Na^+^ diet. Body weight plateaued by about day 3, and in γ^WT/WT^ males, the mean normalized daily body weight from days 3 through 10 was 0.98 ± 0.02, as compared to 0.97 ± 0.02 in γ^Q4/Q4^ males (p = NS). In females, the mean normalized daily body weight from days 3 through 10 was 0.95 ± 0.04 in γ^WT/WT^ and 0.94 ± 0.02 in γ^Q4/Q4^ mice (p = NS). Mixed effects modeling did not reveal any genotype-specific difference in weight as a function of time. Normalized total body water from days 3 to 10 in γ^WT/WT^ males remained stable at 1.01 ± 0.02, compared with 0.98 ± 0.04 in γ^Q4/Q4^ males (p = NS). Over this period in γ^WT/WT^ females, mean daily normalized total body water declined to 0.96 ± 0.02 as compared to 0.99 ± 0.03 in γ^Q4/Q4^ females (p = NS). Mixed effects modeling revealed a significant day – genotype interaction, for male (p < 0.05) but not female (p = NS) mice, suggesting that male, but not female, γ^Q4/Q4^ mice lost body water more rapidly than γ^WT/WT^ controls.

**Figure 3.**
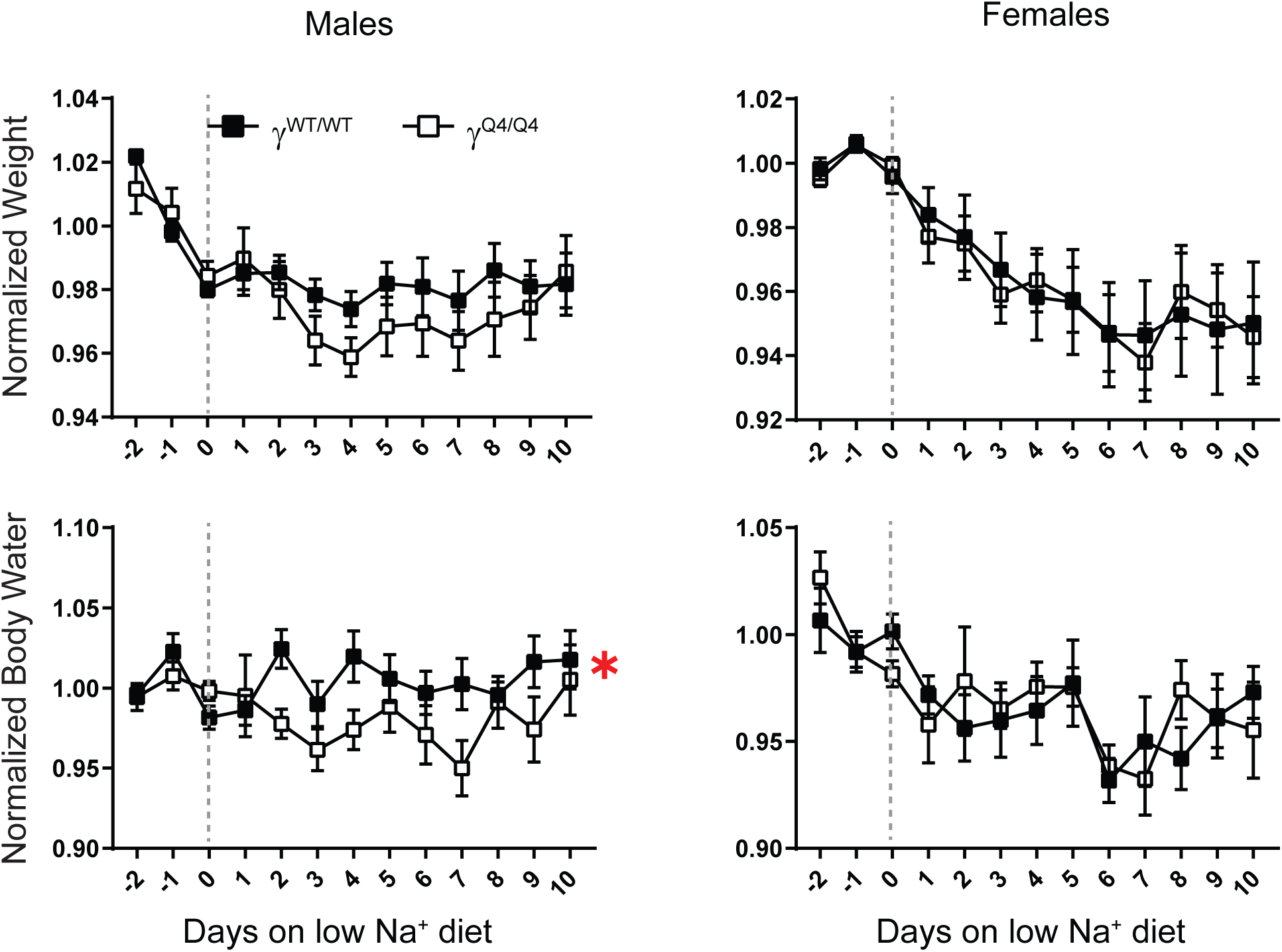
Changes in body fluid and weight in response to a low Na^+^ diet. Normalized body weights (top row) and normalized body water content (bottom row) are shown in male (left column) and female (right column) mice given a low Na^+^ diet for the number of days indicated on the X-axis. Days before 0 represent days on standard (0.23% Na^+^) mouse chow. Error bars represent standard error for N = 6 male γ^WT/WT^, 6 male γ^Q4/Q4^, 7 female γ^WT/WT^ and 8 female γ^Q4/Q4^ mice. In both males and females, change in body weight in response to a low Na^+^ diet was not significantly different in γ^WT/WT^ as compared to γ^Q4/Q4^ mice. In male γ^Q4/Q4^ mice, total body water over 10 days on the low Na^+^ diet was lower than in male γ^WT/WT^ controls (*: p < 0.05, mixed-effects analysis). In females, there was no difference between genotypes in total body water. For clarity, error bars represent standard error.

In exogenous expression systems, proteolytic removal of the inhibitory tract in ENaC’s γ subunit activates the channel by increasing channel P_O_ (13). Removal of the γ subunit’s inhibitory tract requires cleavage at sites both proximal to, and distal to, the inhibitory tract (**Figure 1**). We examined whether disruption of the γ subunit’s furin-cleavage site reduces P_O_ of the channel *in vivo*. Single channel patch clamp recording was performed on microdissected CNT/CCDs from γ^WT/WT^ and γ^Q4/Q4^ mice on a low Na^+^ diet. A channel with ∼8 pS conductance was observed with Li^+^ as the charge carrier (**Figure 4**), consistent with ENaC (64). The number of channels (N) was not significantly different in patches from γ^WT/WT^ (1.8 ± 0.4 channels/patch, N = 5), as compared to γ^Q4/Q4^ (2.8 ± 1.9 channels/patch, N = 8) mice (*p* = NS). Open probability (P_O_) in γ^WT/WT^ (0.05 ± 0.03, N = 5) and γ^Q4/Q4^ (0.08 ± 0.05, N = 7) mice was also similar (*p* = NS). NP_O_ did not differ in patches from γ^WT/WT^ (0.10 ± 0.06, N = 5) and γ^Q4/Q4^ (0.21 ± 0.24, N = 7) mice (*p* = NS). We also observed a previously described 20 pS channel of unclear identity (55). This channel is non-cation selective and is inhibited by high (50 μM) concentrations of benzamil (55). The number of 20 pS channels per patch was similar in γ^WT/WT^ and γ^Q4/Q4^ kidneys (2.1 ± 1.5 channels/patch in γ^WT/WT^ kidneys, N = 16; vs. 2.3 ± 1.4 channels/patch in γ^Q4/Q4^ kidneys, N = 8; *p* = NS), as were the P_O_ (0.47 ± 0.36, N = 16 in γ^WT/WT^ kidneys; 0.49 ± 0.33, N = 8 in γ^Q4/Q4^ kidneys; *p* = NS) and NP_O_ (0.96 ± 0.83 in γ^WT/WT^ kidneys, N = 16; 1.09 ± 0.80 in γ^Q4/Q4^ kidneys, N = 8; *p* = NS). Thus, surprisingly, patch clamp electrophysiology provided no evidence that *in vivo* disruption of the furin cleavage site in ENaC’s γ subunit altered channel activity.

**Figure 4.**
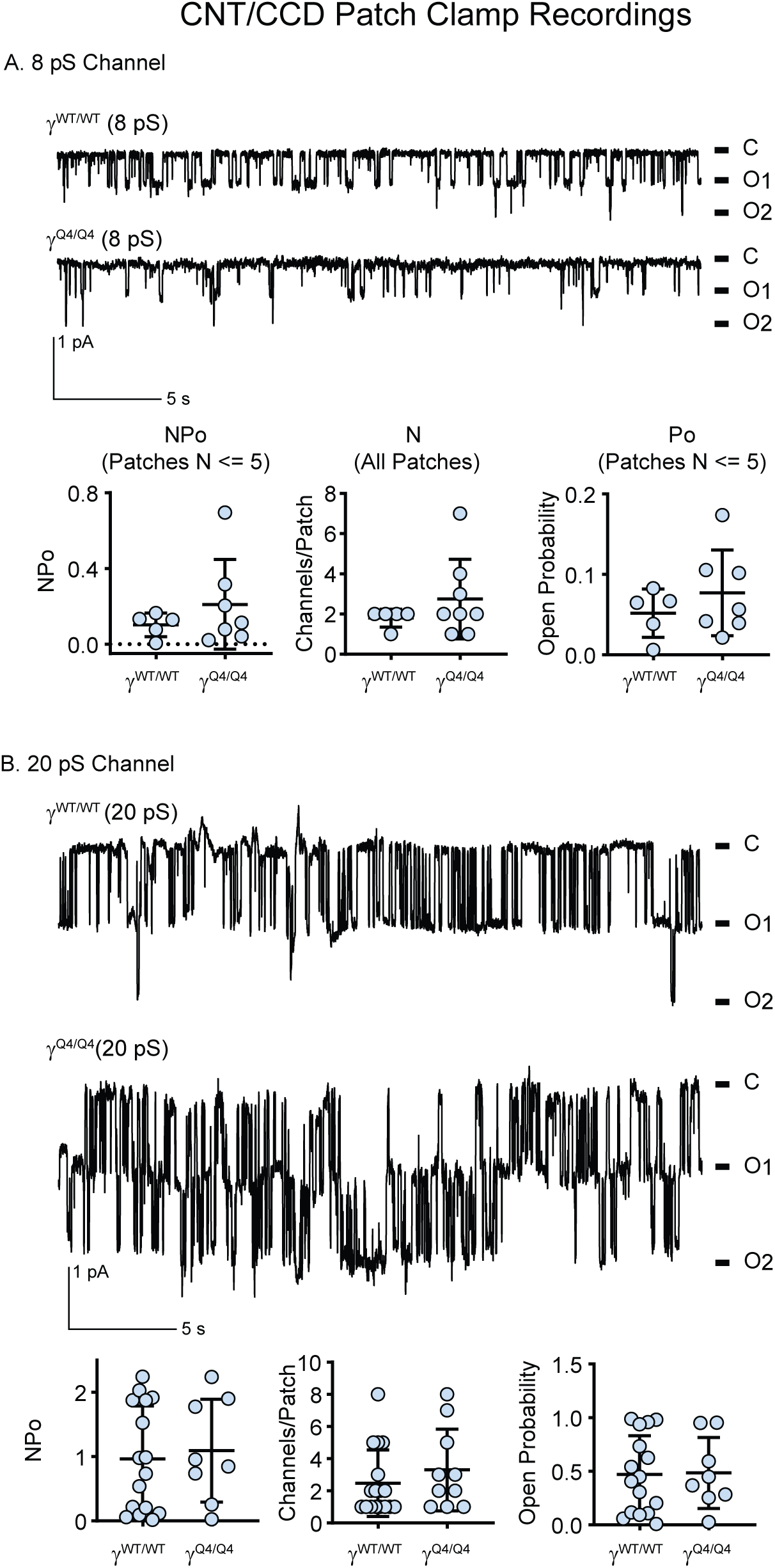
On-cell patch clamp of CNTs/CCDs dissected from male γ^WT/WT^ and γ^Q4/Q4^ mice on a low Na^+^ diet reveal no significant differences in P_O_, channels per patch (N), or NP_O_. A: Representative on-cell current tracings of 8 pS ENaC recorded from dissected tubules from a N = 5 γ^WT/WT^ or N = 7 γ^Q4/Q4^ mice. C represents the closed state. O1 and O2 represent currents when one or two channels are open, respectively. Pairwise comparisons of NP_O_, P_O_, and N (below) show no significant differences in γ^WT/WT^ or γ^Q4/Q4^ mice (p = NS, *t*-test). B: Current tracings of the 20 pS channel. Pairwise comparisons of NP_O_, P_O_, and N (below) show no significant differences in 16 γ^WT/WT^ compared with 8 γ^Q4/Q4^ mice (p = NS, *t*-test). Error bars represent standard deviation of the mean.

We next examined whether collecting ducts from γ^Q4/Q4^ mice on a low Na^+^ diet exhibit altered electrolyte transport in isolated, microperfused tubules. Net Na^+^ absorption (denoted as a positive flux, J_Na_) is stimulated by increased luminal fluid flow rate (58, 65-67). Therefore, we examined J_Na_ and net K^+^ secretion (denoted as a negative flux, J_K_) at a low flow rate (∼1.0 ± 0.1 nL min^-1^ mm^-1^) and a high flow rate (∼5.0 ± 0.2 nL min^-1^ mm^-1^) (**Figure 5**). We predicted that collecting ducts from γ^Q4/Q4^ mice would show a reduced flow-stimulated increase in J_Na_ as compared with collecting ducts from γ^WT/WT^ mice. At a low flow rate, J_Na_ did not differ in collecting ducts from γ^WT/WT^ (10.2 ± 9.2 pmol min^-1^ mm^-1^, N = 5;) and γ^Q4/Q4^ (15.7 ± 4.2 pmol min^-1^ mm^-1^, N = 4, p = NS) mice. Increased flow rate stimulated J_Na_ to a similar degree in γ^WT/WT^ (50.4 ± 6.4 pmol min^-1^ mm^-1^, N = 5) and γ^Q4/Q4^ (55.9 ± 16.7 pmol min^-1^ mm^-1^, N = 4; p = NS) collecting ducts.

**Figure 5.**
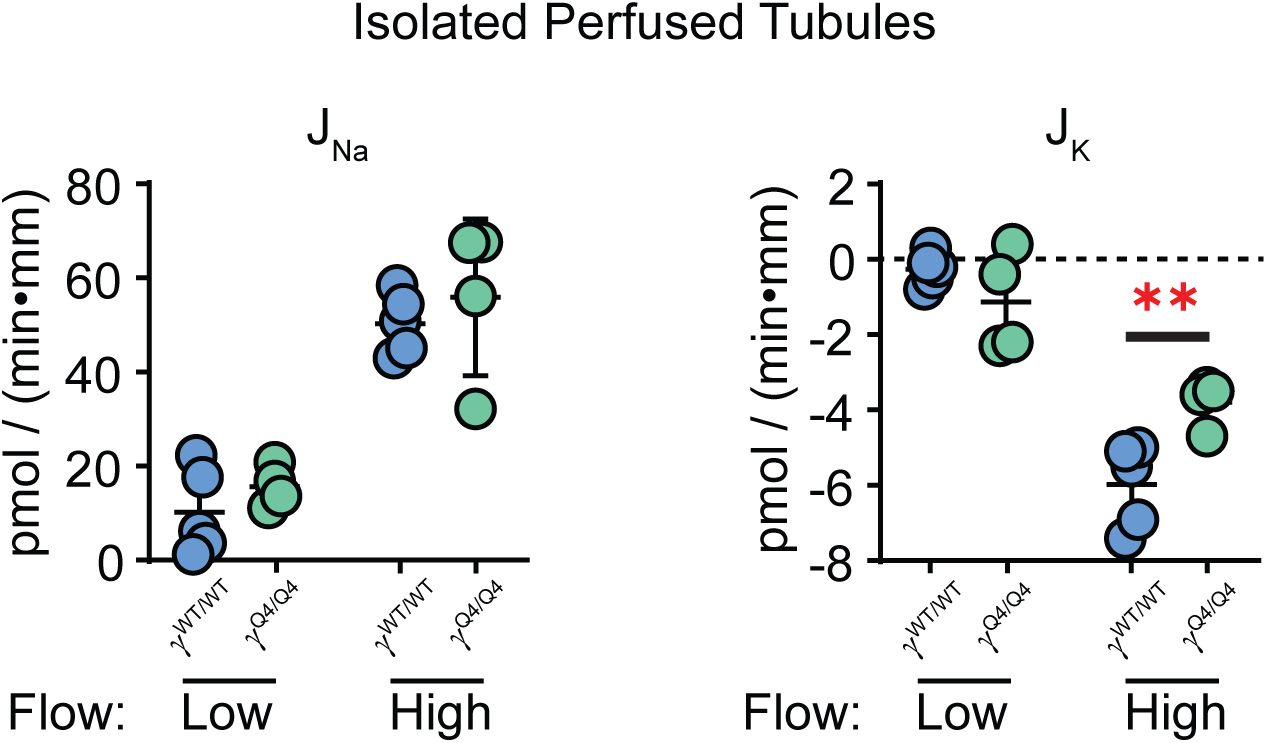
Measurements of J_Na_ and J_K_ in isolated perfused cortical collecting tubules. Tubules were dissected from male mice on a low Na^+^ diet. Luminal flow rates were either low (1 nl/min) or high (5 nl/min). J_Na_ was not different on the basis of mouse genotype at either low or high flow rates. At a low fluid flow rate, J_K_ was similar in γ^Q4/Q4^ and γ^WT/WT^ mice (N=5 γ^WT/WT^, and 4 γ^Q4/Q4^ mice. p = NS, *t*-test). However, at a high flow rate, J_K_ was significantly higher in tubules from γ^WT/WT^ than γ^Q4/Q4^ mice (N=5 γ^WT/WT^, and 4 γ^Q4/Q4^ mice. **: p < 0.01, *t*-test). Error bars represent standard deviation of the mean.

Increased luminal flow also stimulates J_K_. Tubular K^+^ secretion is inhibited by ENaC blockade, demonstrating its dependence on ENaC activity (68, 69). We predicted that collecting ducts from γ^Q4/Q4^ mice would exhibit reduced stimulation of J_K_ at high flow rate, compared with γ^WT/WT^ mice. At low flow, J_K_ was similar in collecting ducts from γ^WT/WT^ (-0.3 ± 0.4 pmol min^-1^ mm^-1^, N = 5) and γ^Q4/Q4^ (-1.1 ± 1.3 pmol min^-1^ mm^-1^, N = 4; p = NS) mice. High flow increased the magnitude of J_K_ in collecting ducts from both γ^WT/WT^ (-6.0 ± 1.1 pmol min^-1^ mm^-1^, N = 5) and γ^Q4/Q4^ mice (-3.8 ± 0.6 pmol min^-1^ mm^-1^ collecting ducts, N = 4). The flow-induced increase in K^+^ secretion in collecting ducts from γ^WT/WT^ mice was significantly greater than in collecting ducts from γ^Q4/Q4^ mice (p < 0.01).

ENaC is expressed in mouse distal colon, and its activity is markedly enhanced when mice are placed on placed on a low Na^+^ diet. Distal colons were harvested from mice on a low Na^+^ diet for 14 days and placed in Ussing chambers bathed in Ringer’s solution. Short-circuit current (I_SC_) was monitored before and after the addition of 100 μM amiloride to the apical bath. Representative I_SC_ recordings are shown in **Figure 6**. Amiloride-sensitive I_SC_ was similar in colons from γ^WT/WT^ and γ^Q4/Q4^ mice (-120 ± 101 μA/cm^2^ in γ^WT/WT^ mice, N=6; -198 ± 58 μA/cm^2^ in γ^Q4/Q4^ mice, N=6; p=NS)

**Figure 6.**
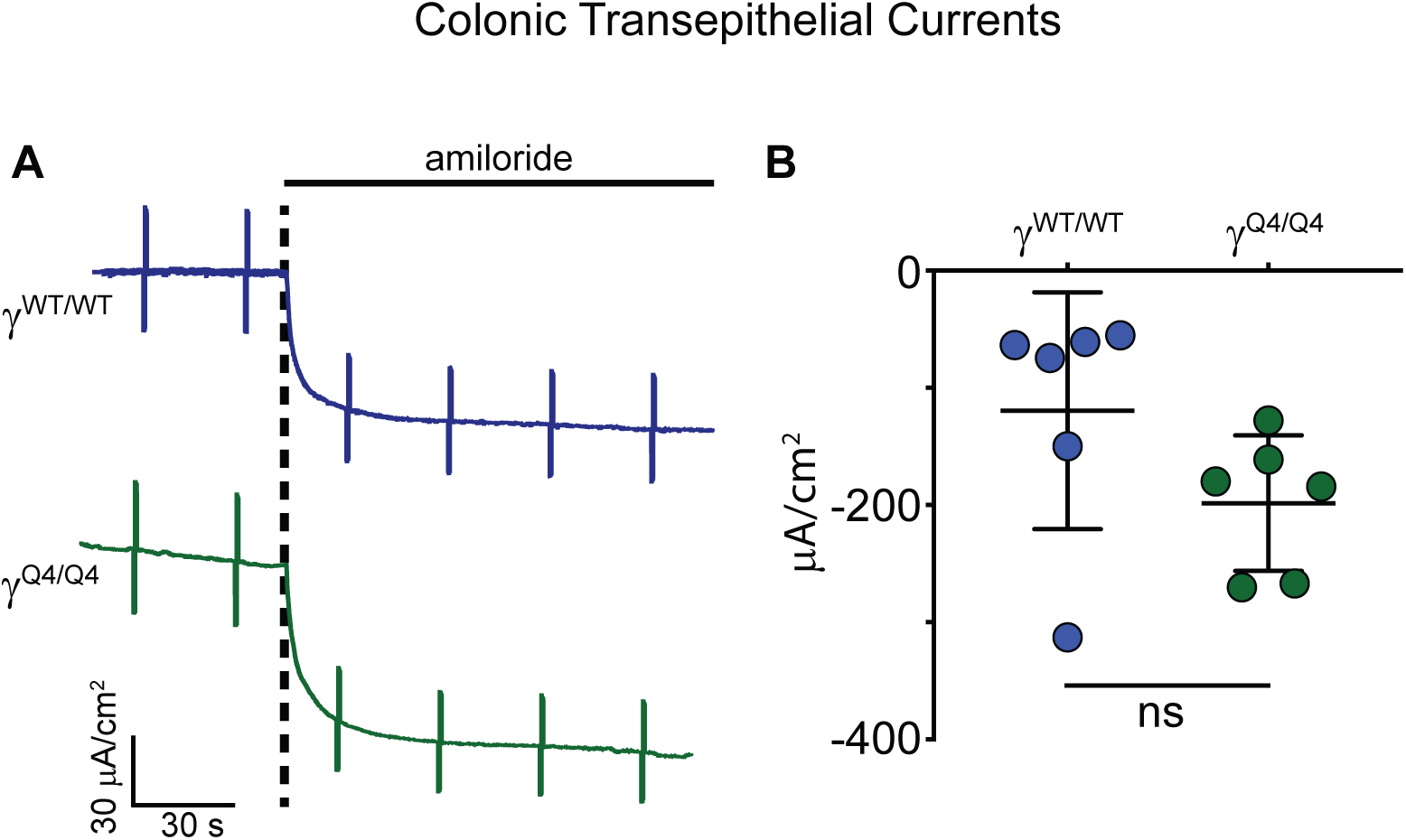
Amiloride-sensitive currents in colonic epithelium. A: Representative short circuit current tracings from colonic epithelia dissected from male γ^WT/WT^ and γ^Q4/Q4^ mice are shown. B: Quantitative summary of amiloride-sensitive current amplitudes shows no significant difference in colonic epithelia from N=6 γ^WT/WT^ compared with N=6 γ^Q4/Q4^ mice (p = NS, unpaired t-test). Error bars represent standard deviation of the mean.

## DISCUSSION

This study confirmed that interruption of the mouse ENaC γ subunit furin cleavage site prevents activation of ENaC by the serine protease prostasin in channels expressed in *Xenopus* oocytes, as previously reported (12). It also demonstrated that γ^Q4/Q4^ mice lacking the γ subunit furin cleavage site exhibit loss of the 57 kDa γ subunit C-terminal fragment noted following PNGase F treatment, which corresponds to the C-terminal γ subunit furin cleavage fragment. These results confirmed that the γ subunit in γ^Q4/Q4^ mice is resistant to furin-mediated proteolysis. A band migrating slightly slower than 57 kDa was noted in male kidney lysates from both WT (without PNGase F treatment) and γ^Q4/Q4^ mice (with and without PNGase F treatment), suggesting that this is a non-specific band. Although we included this band in our analysis of the 57 kDa band intensity in PNGase F-treated lysates from the γ^Q4/Q4^ mice (Figure 2), analyses of the immunoblots excluding this band suggested there was virtually no γ subunit furin cleavage in kidneys from males and female γ^Q4/Q4^ mice. Given these observations and the dramatic difference in proteolytic activation of the channel *in vitro*, we expected to see large and significant phenotypic differences between WT and γ^Q4/Q4^ mice, but this was not what we observed.

The γ^Q4/Q4^ mice were viable and appeared healthy. At baseline, these mice exhibited no difference in body weight, body fluid, or plasma electrolytes. A high K^+^ diet uncovered no difference in whole blood electrolyte values, with the exception of a higher plasma Cl^-^ in blood from γ^Q4/Q4^ males on a high K^+^ diet. On a low Na^+^ diet, no differences in plasma electrolytes or aldosterone levels were evident. Although ENaC is expressed in the colonic epithelium, amiloride-sensitive short-circuit currents in colonic epithelia from γ^Q4/Q4^ and γ^WT/WT^ mice a low Na^+^ diet were similar.

In isolated, microperfused tubules from male γ^Q4/Q4^ mice on a low Na^+^ diet, no difference in flow-induced stimulation of J_Na_ was observed, compared with γ^WT/WT^ controls. However, tubules from γ^Q4/Q4^ mice did exhibit attenuated flow-induced stimulation of J_K_. We recently demonstrated that BK channels in intercalated cells mediate flow-induced K^+^ secretion (58). The reduced flow-induced J_K_ seen in tubules from γ^Q4/Q4^ mice suggests an altered lumenal potential, even though differences in J_Na_ were not observed.

Perhaps the most perplexing findings were that a loss of the γ subunit furin cleavage site did not demonstrably change either activity of the 8 pS channel (measured with Li^+^ as the charge carrier) or J_Na_ in isolated tubules. In exogenous expression systems, proteolytic processing clearly activates ENaC, increasing P_O_ without changing unitary conductance (70-72). It is notable that the P_O_ of the 8 pS channels on a low Na^+^ diet was low. This likely reflects our use of 129/Sv as a background strain and is consistent with our other ENaC single channel recordings in CNTs/CCDs from this mouse strain (55). Recent evidence showed that impaired ENaC activity due to a γ subunit mutation that prevents its palmitoylation was associated with increased activity of a higher conductance (20 pS), non-selective cation channel that is blocked by a high (50 μM) concentration of benzamil (55). We determined NP_O_, N and NP_O_ of the 20 pS channel in CNTs/CCDs from WT and γ^Q4/Q4^ mice on a low Na^+^ diet. We also detected no differences in the activity of this higher conductance channel. The similar ENaC P_O_ we observed in CNTs/CCDs from WT and γ^Q4/Q4^ mice agrees with the similar levels of amiloride-sensitive Na^+^ currents that we observed in distal colonic epithelia from either γ^Q4/Q4^ or WT mice maintained on a low Na^+^ diet.

Despite that lack of a difference in ENaC P_O_, we observed a small, but significant, impairment in body fluid conservation in the context of dietary Na^+^-restriction in male γ^Q4/Q4^ mice, when compared to WT. This observation suggests that there may be modest differences in ENaC activity that were we unable to discern in our patch clamp or microperfused tubule studies. Previous studies suggested that factors associated with enhanced proteolytic processing of ENaC subunits also lead to an increase in Na^+^ reabsorption in the aldosterone-sensitive distal nephron. Examples of this include dietary Na^+^-deprivation (9), aldosterone infusion (10), and direct tubular application of proteases (7). Conversely, pharmacologic inhibition of proteolysis in the kidney enhances Na^+^ excretion (73-75). We did not observe impaired body fluid conservation in female γ^Q4/Q4^ mice on a low Na^+^ diet. Previous reports described similar ENaC expression and γ subunit proteolytic processing in male vs. female mice (76), and we do not have an explanation for these sex-specific differences. It is possible that other experimental perturbations capable of stimulating ENaC subunit proteolysis, renal Na^+^ retention, and volume expansion (i.e., nephrotic syndrome and aldosterone administration) could result in phenotypic differences between WT and γ^Q4/Q4^ mice.

After more than 40 years of studies exploring the influence of proteases on ENaC function and Na^+^ handling, this report represents the first study examining the *in vivo* importance of furin-mediated proteolysis of ENaC’s γ subunit upon bodily Na^+^ and K^+^ homeostasis. Despite the large effects on ENaC activity that have been described in association with γ subunit proteolytic processing and release of its inhibitory tract *in vitro*, the effects that we observe *in vivo* on ENaC activity (i.e., channel P_O_), transepithelial Na^+^ transport in CCDs and distal colon, and volume regulation were modest, at best. Our results suggest that other factors have important roles in modulating ENaC activity and compensate for a lack of γ subunit furin processing in the γ^Q4/Q4^ mice placed on a low Na^+^ or high K^+^ diet. These findings are consistent with a recent report showing that deletion of the gene encoding the serine protease prostasin, a known activator of ENaC (see Figure 1), in kidney tubules was not associated with reduced ENaC function (77). Numerous reports in rodents use γ subunit proteolysis as a marker of ENaC activation. While we agree that assessing the extent of ENaC subunit proteolysis is important, our work suggests that proteolytic processing should not be relied upon in isolation as a primary indicator of *in vivo* ENaC activity.

## AUTHOR CONTRIBUTIONS

E.C.R., S.S., S.G., A.N., T.R.K. conceived and designed research.

E.C.R., S.S., L.Z., C.B., Z.K., S.G., A.N., T.L., A.L., A.W., A.J., R.C. performed experiments.

E.C.R., S.S., S.G., A.N., A.J., R.C., L.M.S., T.R.K. analyzed and interpreted data.

E.C.R., S.S., A.N. prepared figures.

E.C.R. drafted manuscript.

E.C.R., A.N., R.C., L.M.S., A.K., T.R.K. edited and revised manuscript.

All authors approved the final version of this manuscript.

